# A novel cost framework reveals evidence for competitive selection in the evolution of complex traits during plant domestication

**DOI:** 10.1101/2020.10.30.362236

**Authors:** Robin G Allaby, Chris J Stevens, Dorian Q Fuller

## Abstract

Most models of selection incorporate some notion of environmental degradation where the majority of the population becomes less fit with respect to a character resulting in pressure to adapt. Such models have been variously associated with an adaptation cost, the substitution load. Conversely, adaptative mutations that represent an improvement in fitness in the absence of environmental change have generally been assumed to be associated with negligible cost. However, such adaptations could represent a competitive advantage that diminishes resource availability for others and so induces a cost. This type of adaptation in the form of seedling competition has been suggested as a mechanism for increases in seed size during domestication, a trait associated with the standard stabilizing selection model. We present a novel cost framework for competitive selection that demonstrates significant differences in behaviour to environmental based selection in intensity, intensity over time and directly contrasts to the expectations of the standard model. Grain metrics of nine archaeological crops fit a mixed model in which episodes of competitive selection often emerge from shifting optimum episodes of stabilizing selection, highlighting the potential prevalence of the mechanism outlined here and providing a fundamental insight into the factors driving domestication.

---

Natural selection that gives rise to the differential reproductive success of competing individuals of varying fitness leads to the notion first pointed out by Haldane (1957) of a cost in terms of a declining population size during adaptation. Such costs lead to theoretical selection limits which have been extensively debated in terms of their severity, depending in part on pleiotropic effects between alleles, alternative adaptive solutions and modes of selection such as truncation selection in selective breeding (Barton 1995, Maynard-Smith 1968, Sved 1968). Regardless of the incorporation or not of cost penalties, most models of selection work on a similar basis to that originally described by Haldane in which individuals of a population are universally disadvantaged in regard to some parameter, from which fitter individuals evolve. In other words a universal genetic load is bestowed on the population because of some change in their environment. A second category of criticism to the original Haldane model was that such universal genetic load may not represent the majority of adaptive evolution. Instead, selection may favour ‘advantageous’ alleles in the absence of any environmental degradation or punishment of the wild type (Brues 1964, 1969), and could represent improvements such as in size (Van Valen 1963). Such selective episodes were thought unlikely to carry a cost largely because of the absence of a genetic load (Brues 1964,1969, Van Valen 1963, Felsenstein 1971), if the population were permitted to expand.

However, if a population has reached the current carrying capacity of its environment, as is expected under a niche concept, and therefore has no capacity to expand, then a slight cost is expected to be shared among the wild type population as a competitive pressure is applied from the adapted phenotype. Intuitively, it is implied that as the adapted phenotype increases in frequency, the relative disadvantage experienced by the wild type population will increase and an intensification of selection is predicted. Conceptually, this contrasts to the environmental degradation model of selection as rather than a sudden appearance of a large genetic load across the population determined by some environmental factor a small genetic load is generated by the appearance of an innovative mutation, which then will grow in effect over time as the mutation gains frequency.

This category of selection may apply, for instance, in the case of increased seed size during plant domestication where it has been argued that seedling competition may be responsible selecting for more robust early growth (Fuller and Stevens 2017, 2019). An increased ability for resource acquisition, in this case nutrients and light, suggests a cost to the remaining population when resource is finite. Seed size is a complex trait influenced by many loci of varying effect which are well described under the standard stabilizing selection model (Barton 1986). This latter represents a category of environmental selection models in which the strength of selection decreases as the optimum is approached and pleiotropic constraints apply. While much is understood about the potential interactions of alleles of varying effect from numerous different loci in the process of adaptation through simulations (Chevin *et al*. 2008, Jain and Stephan 2017, Pavlidis *et al*. 2012, Thornton 2019), here we consider the fate of a single allele of varying effect on a competitive trait advantage and then attempt to place that within the wider context of complex trait adaptation.

## METHODS and MODELS

### Basic model rationale and validation

In this model we assume that fitness is associated with a probability of acquiring resource. Broadly, this may be any organism that first encounters a resource and then does, or does not, acquire it. We assume that individuals have equal encounter probabilities but differ principally in their ability to acquire the resource once encountered. The probability of an adaptive mutant and wild type acquiring a resource on encounter are given is *p_m_* and *p_w_* respectively. Consequently, the probability of a resource being acquired (*P(aq)*) on encounter can be given as:

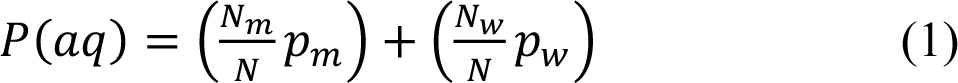

Where *N_m_, N_w_* and *N* are the number of mutant phenotypes, wild types and total population size respectively. If we assume that the resource is finite and is exploited completely, then the expected number of encounters, or trials (*tr*), required before each unit of resource (*r*) is consumed can be given as:

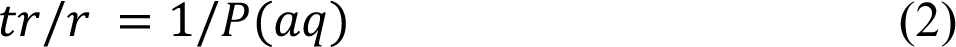

The parameter *tr* will follow a Poisson distribution with λ = *tr*, but here we take the λ value as a simplifying estimate adequate for the purposes of the model. Consequently, the expected resource (*E(m)*) a single mutant phenotype will obtain can be given as the product of probability of a selected mutant phenotype for a given trial, the probability the individual will acquire the resource after encounter, and the total number of trials:

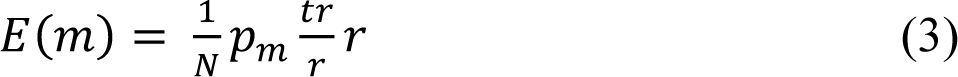

We can measure the selective advantage of a mutant by considering the increased resource acquisition relative to the equal share that would be received as expected under no advantage. The excess, or advantage (*a*) that the mutant will obtain relative to the wild type can be given as the increased proportion of resource relative to equal share:

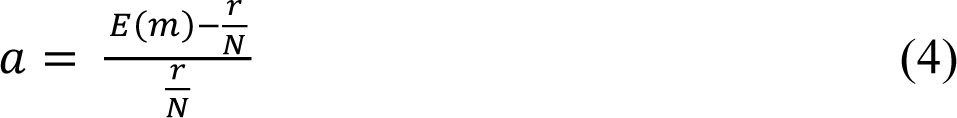

The remaining wild type population will be relatively disadvantaged by an amount that is shared equally amongst them, which will reduce the fitness of the wild type relative to the adaptive mutant. If we assume that a niche is maximally exploited, it seems a reasonable assumption that an equal share represents the sufficient amount required for survival. The reduced proportion relative to an equal share available to the wild type can be taken as the probability of obtaining sufficient resource. Under this assumption we can take the reduction in probability of sufficient resource acquisition as the selection coefficient (*s*). Equation (3) describes the diminishing resource adaptive mutants acquire as they increase in frequency and consequently the total number of trials required to utilize total resource reduces. For any frequency of adaptive mutant individuals the extra resource acquired per individual will have an additive effect on the remaining wild type population which share the burden of reduced resource such that the selection coefficient experienced by any single wild type individual can be expressed as:

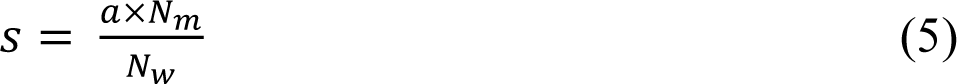

Equation (5) expanded using equations (1-4) and simplified, with *r* assigned a value of 1, express the selection coefficient as:

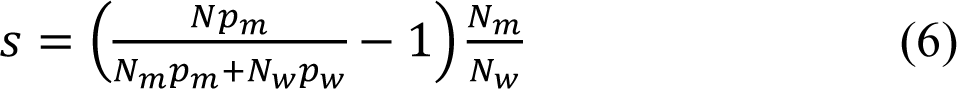

equation (6) is validated by an agent-based simulation in which mutant and wild type agents are assigned respective values of *p_m_* and *p_w_*, and the resulting proportional loss of resource relative to an equal share measured, Fig. S1. The behaviour of parameters as described in 6quation (6) was verified using a simple agent-based simulation in which individuals were randomly selected and then deemed successful in resource acquisition with a probability of *p_m_* or *p_w_* depending on whether they were wild type or mutant individuals, Fig. S1. Sampling continued until all resource had been allocated. Arbitrary values of 0.21 and 0.2 were assigned to *p_m_* and *p_w_* respectively in a population size of 100 individuals with resource size 1000 and 10,000 replications. Sampling was repeated for increasing numbers of mutants from 1-99. The perl script used for this (benign_profile_of_s.pl) is available from (https://warwick.ac.uk/fac/sci/lifesci/research/archaeobotany/downloads/competitive_selection).

This validation demonstrates that the potential effects of increasing competition between mutants as their frequency increases does not change the outcome. Consequently, despite the ‘zero sum game’ for adaptive mutants in that their advantage becomes less over time until at the point of allele fixation there is no advantage, the selection system works with increasing and predictable intensity because of the increasing cost over time to the wild type. We can therefore apply the rules of equation (6) to ascribe idealized selection coefficients within a competitive selection model.

A remaining apparent problem is that we do not necessarily know the values of the variables *p_m_* and *p_w_*. equation (6) can be rearranged for the ratio of *p_m_* and *p_w_* for a single mutant:

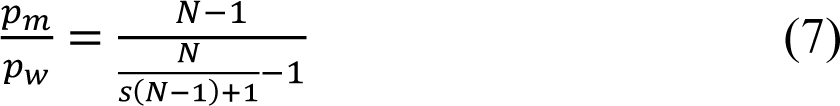

Given that the intrinsic success probabilities on encounter are constant, the ratio holds true for any number of *N_m_* and *N_w_*, and consequently the ratio itself is dependent only on the population size and the selection coefficient. A corollary is that any two probabilities for *p_m_* and *p_w_* that satisfy the ratio defined by *s* and *N* can be used to calculate *s* in equation (6).

Therefore, any probability can be assigned to one variable (*p_m_* or *p_w_*) and equation (7) can be used to calculate the corresponding variable and we do not need to know the true values of *p_m_* and *p_w_* to find *s*, only a pair of values that satisfy equation (6) for a particular value of *s*.

### Competitive selection simulations

A previously described environmental selection agent-based simulator (Allaby *et al*. 2015) was developed to include a competitive selection regime using the equations presented here. Briefly, the model simulates a population of diploid individuals of defined size and number of loci under selection that are assumed unlinked. A single adaptive mutation is seeded into the population for each locus, and reseeded if lost through drift. Each generation new individuals are generated by random gamete union, and individual survival is determined stochastically with a probability equal to the fitness for each locus under selection.

Consequently, selection is against the wild type. Population generation is limited by a defined fecundity parameter, resulting in diminishing population sizes when the substitution load is high. The modification of the model for this study was that each generation frequency dependent selection coefficients against the wild type were calculated for a given number of advantageous phenotypes using equation (6), deriving values of *p_m_* or *p_w_* using equation (7). The simulator is available from (https://warwick.ac.uk/fac/sci/lifesci/research/archaeobotany/downloads/competitive_selection).

### Threshold frequency determination

Although it is well known that selection proceeds at a pace independent of population size in contrast to neutral drift, at the initial low frequencies of adaptive mutants, drift can be stronger than selection. We considered a potential threshold at which the expected increase in frequency of an allele *p* under selection (δ*p*) exceeds the proportion of a single individual allele in the population (1/2*N*). The value of δ*p* for a given inbreeding coefficient (*F*) is given as:

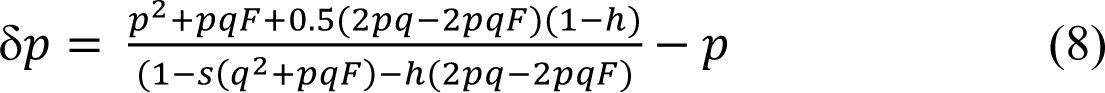

Where *h* is the selection coefficient against the heterozygote, *q* = 1-*p* and *s* can be calculated from equation (6). For a given number of mutant phenotypes (*N_m_*) the frequency of the mutant allele *f(r)* was determined following the solution to the quadratic equation derived from Hardy Weinberg for recessive mutants (where *N_m_/N* = *q^2^ + pqF*) as:

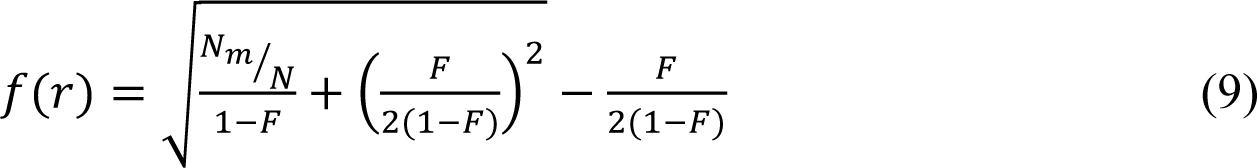

And for dominant mutants *f(d)*, where *N_m_/N* = *1-( q^2^ + pqF)*:

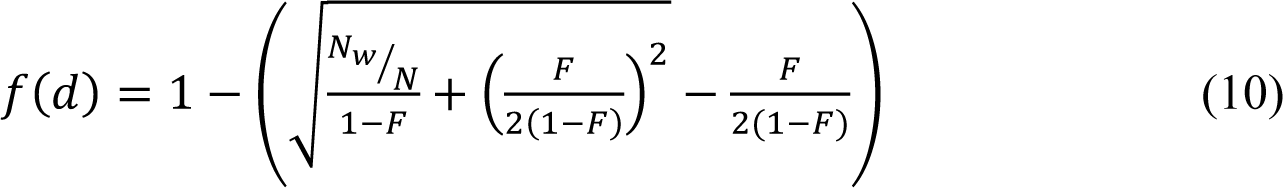

For each number of mutant phenotypes in the population, equations (9) or (10) were used to determine the frequency of *p*, from which the frequency dependent value of *s* was determined using equation (6), and then equation (8) used to determine δ*p*. Once the frequency of *p* is determined at which δ*p* > 1/2N, the neutral probability of an allele reaching that frequency is given as 1/*f_(threshold)_*, where *f_(threshold)_* is the frequency of *p* at which δ*p* > 1/2N. Similar thresholds were calculated for environmental selection which differs only in that *s* is constant against wild types rather than frequency dependent.

### Distribution of fitness effects for competitive and environmental selection

The probability of various ratios of *p_w_* and *p_m_* occurring is expected to be described in the distribution of fitness effects, which are well described by an exponential distribution (Gillespie 1983, 1984, Orr 2003). The probability of the magnitude in difference between *p_w_* and *p_m_* can therefore take the form:

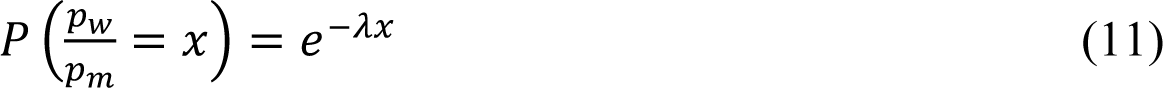

In which the parameter λ takes a value of 1 as a general solution (Orr 2003). Similarly, we used equation 2 of Orr (2003) to describe the distribution of fitness effects of *s* for environmental selection.

### Relative likelihood profiles of selection intensity

We calculated how likely different selection intensities of competitive and environmental selection were to occur respectively by considering 1. the probability of selection strength given a distribution of fitness effects, 2. the probability that mutant alleles would reach sufficiently high frequencies through drift at which δp > 1/2N and so selection would become dominant over drift, 3. the mutation rate, and also 4. the probability of population survival for a given magnitude of selection as previously calculated (Allaby *et al*. 2015). The latter parameter only came in to effect for *s* equal or greater than 0.5 and was otherwise 1 for all other instances. Rate of mutation generation (2*Nµ*) expresses the importance of *N* rather than *µ*, the latter was therefore only notionally included here with *µ*=0.5 for convenience of display. Hence likelihood profiles are relative for comparison between selection intensities. Likelihood profiles were therefore calculated as the product of these four probabilities.

### Competitive selection profiles

Theoretical competitive selection tracks were calculated for given values of initial *s* and *N* where values of *p* increment each generation by δ*p.* For each new value of *p* each generation values of *N_m_* and *N_w_* were determined using the Hardy Weinberg expectations for a given mating system in mutant dominant allele systems:

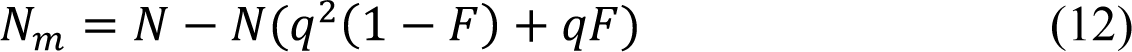

And in the case of recessive advantageous alleles:

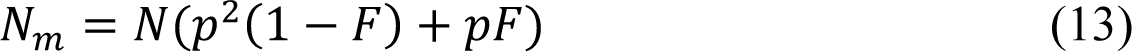

These values were used in equation (6) to determine *s,* which was used to determine δ*p* with equation (8), and consequently the value of *p* in the next generation. The process was iterated through to mutant allele fixation.

### Standard model selection profiles

The standard model of stabilizing selection (Barton 1986) was used to compare predictions of selection coefficient profiles against real data. The standard model assumed the conventional relationship between fitness (1-*s*), size trait values (*z* values) and selection intensity as described by the genetic variance (*V_s_*) such that:

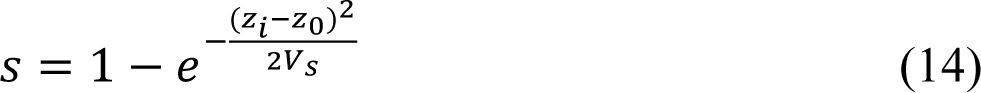

Where *z_0_* is the optimal trait value, and *z_i_* is the *i*th trait value during trait value iteration. Since it is the intensity of selection which determines the rate at which traits change rather than the magnitude of the trait change, an initial value of 1 was assumed for *z*-*z_0_*, such that intensity of selection is given by:

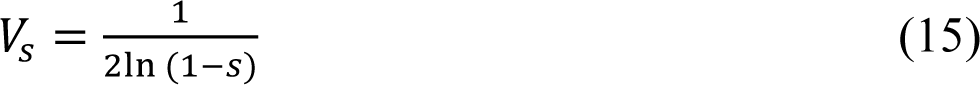

To determine the extent to which a trait changed per generation under intensity *Vs*, *s* was first converted to haldanes:

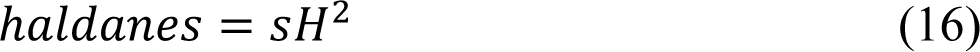

Where *H* is the heritability parameter. The iterate *i*th trait was then determined using a derivation of equation (18):

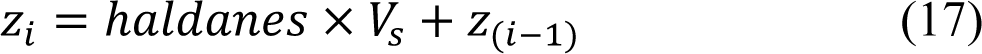

To generate a selection track an initial value of *s* was used firstly calculate *V_s_* (eq 15), haldanes (eq 16), and the extent to which the trait value changed (eq 18). The modified trait value was then used to recalculated *s* (eq 14) and the process was repeated for a constant value of *V_s_* until the difference in optimal and actual trait values was less than 0.0001.

### Calculation of s from archaeometric data

Selection coefficients were determined by first calculating haldanes using the method of Kinnison and Hendry (2001) in which:

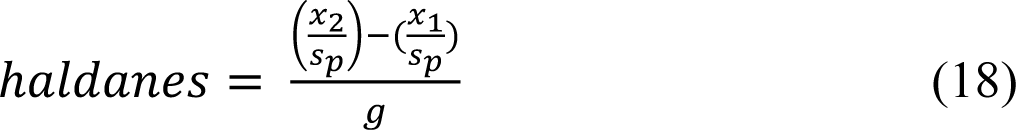

where *x* is the mean trait value at time 1 and 2, *s_p_* is the pooled standard deviation of trait values across time, and *g* is the number of generations which is assumed to equate to radiocarbon date years in the case of annual crops. Haldanes were converted to *s* following

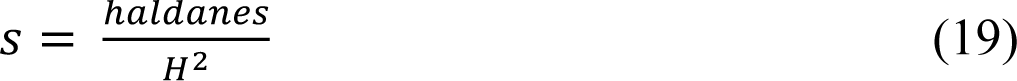

where *H* is the trait heritability, assumed here to be approximately 0.9 for size traits (Fuller *et al*. 2014). All archaeometric measurements of seeds were taken from the supplementary data of Fuller *et al*. (2014). Using this approach, values of *s* were calculated for all data point pairs which were less than 1000 years and more than 100 years apart in age. The exception was in the case of sunflowers, where the low number of useful data points was mitigated by considering data points up to 2000 years apart.

### Model fitting to archaeometric data

Competitive and standard models of selection were fitted to series of real archaeometric estimates of selection coefficients. Selection tracks were iteratively generated for each model as described above until the fit could not be improved. The best fit for each track was achieved using custom scripts by sliding the track along time (*x* axis) and finding the least squares difference in *s* values (*y* axis) at real data points in time. The fit was then evaluated by calculating *r* as a linear regression of y_theoretical_ against y_observed_ coordinates. In the case of competitive selection tracks began with a lowest value of *s* determined as *s_max_*/4*N* where *s_max_* is the highest selection coefficient in the series of archaeometric *s* estimates. Values of *s* were iteratively increased by 0.0000001 until a maxima in *r* was reached. To check for spurious correlation or over fitting of the competitive model we calculated *p_null_*, the probability that observed *r* could occur for a random upward trend. To do this we compared *r* values against null distributions of *r* values generated as follows. Using constraints of archaeometric values of *s* bounded by values of 0.0001 and 0.02 and time bounded by archaeological sample ages (*t*), we randomly generated data sets by first generating two values of *s* and *t* to give upper and lower *s* values of a random upward trend. Intervening data points were then randomly generated which were constrained by the upper and lower values of *s* and *t* initially generated. The number of data points generated matched the observed number of data points. We then fitted a competitive selection curve to the data and calculated the *r* value as described above. For each observed *r* value we generated 1000 such replicates to produce a non-parametric null *r* distribution. The proportion of replicate *r* values above the observed value gave our estimate of *p_null_*.

In the case of the standard model tracks the same sliding approach as described above to find the best fit for each track. Initial values of *s* were incremented by 0.00001 until *r* values were maximized, and each archaeometrically determined value of *s* was examined to give the highest overall *r*.

### Sliding window scans of competitive and standard models

To initially scan through the archaeometric datasets and determine whether the standard or competitive model showed the better fit to the data a sliding windows of 5 or 10 (depending on data density) temporally consecutive selection coefficients were used. For each window the best fitting selection track was determined for each model as described above, and windows were slid by one selection coefficient at a time. In the case of competitive selection sliding scan fits were based on a population size of 1000.

## RESULTS

### Relative intensities of competitive and environmental selection

The additive nature of the strength of competitive selection as mutants increase in frequency as described in equation (6) suggests that the initial selection coefficients involved may be very small and considerably weaker than genetic drift. In such cases selection could not effectively act until mutant allele frequencies had been elevated sufficiently through drift processes. The interplay between drift and selection is largely dependent on the parameter δ*p*, the expected change in frequency of an adaptive mutant due to selection over a single generation. The point at which δ*p* becomes larger than the proportion of a single allele in the population is a likely threshold because below this no increment in *p* is expected in terms of individuals in the population, and the behaviour of *p* will be close to neutral. Consequently, at frequencies of *p* at which δ*p* exceeds 1/2N, it is expected that there should be a shift from more drift-like behaviour to more selection like behaviour to an extent that is determined by the probability that *p’>p*, where *p’* is the value *p* becomes in the subsequent generation.

Simulations under the competitive model broadly support this threshold behaviour showing a biphasic process in which advantageous mutants wander in a drift like fashion at lower frequencies, but switch into strong selection with increasing frequency, Fig. S2. Under an environmental selection model, it is well known that intensities (*Ns*) >> 1 are required for selection to overcome drift. However, despite an initial section intensity of only 0.5 competitive selection is still effective, and remains so at initial intensities ten-fold lower, Fig. S3. Similar biphasic behaviour is of course expected under the standard environmental selection model, although at such low selection intensity the influence of drift remains great because selection does not intensify with increasing frequency of adaptive mutants, and consequently the probability that *p’* will be higher than *p* remains marginal, Fig. S4.

Together, these results suggest that the probability of mutations reaching δ*p* > 1/2N threshold is a reasonable approximation to describe the probability of the successful selection of an adaptive mutant.

The level of the drift threshold, and the strength of selection that occurs once an adaptive mutant has broken through it is largely influenced by the fitness effect of the mutation. While in the case of environmental selection the fitness effect is described by the selection coefficient *s*, in competitive selection the fitness effect is described by the *p_w_/p_m_* ratio and *s* is a function of the ratio and the frequency of the mutant. The probability of the magnitude of difference between values of *p_w_* and *p_m_* is therefore expected to follow a distribution of fitness effects, which are likely to be well described by an exponential distribution (Gillespie 1983, 1984, Orr 2003). Unlike *s*, the *p_w_/p_m_* ratio has the property of being independent of *N* for a given selection intensity (*Ns*), Fig. S5. Therefore, a single *p_w_/p_m_* ratio describes all population sizes undergoing competitive selection for a particular intensity. Consequently, a probable fate of a competitive mutant can be described as an interplay between the *p_w_/p_m_* ratio and the neutral probability of reaching threshold frequencies at which selection is sufficient to increase mutant frequencies by an individual or more each generation, Fig. S6. This demonstrates that in order for a high probability of fixation *p_m_* must be considerably higher than *p_w_* to achieve the necessary intensity of selection, and the larger the population size the more pronounced must be the difference between the *p_m_* and *p_w_* probabilities. An unsurprising corollary is that competitive mutants are generally more likely to be successful in small populations.

Consequently, by considering for a given selection intensity the rate of mutation generation at 2*Nµ*, the probability of fitness effect, the probability of reaching selection/drift threshold frequencies and accounting for the probability of population survival of substitution loads (Allaby *et al*. 2015), we can produce relativistic likelihood profiles for selection intensities for competitive and environmental selection respectively, Fig.1. Our first major finding is a clear separation of the most likely initial selection intensities under competitive versus environmental based selection. Whereas we find that environmental based selection is expected to occur at intensities >> 1 as expected, and the optimal value for *s* falls around 0.05 in agreement with estimates from variance analysis based on empirical data (Johnson and Barton 2005), competitive selection intensities are most likely in this model at values around 0.5. These results suggest that the most likely *p_w_/p_m_* values to successfully undergo selection are around 0.75 or that mutants that are 50% more likely to acquire a resource than the wild type are most likely to occur. This suggests alleles of typically large effect are most likely to undergo selection in this way, although effects could commonly range from 1% to 200% in increased likelihood of resource acquisition.

**Figure 1.**
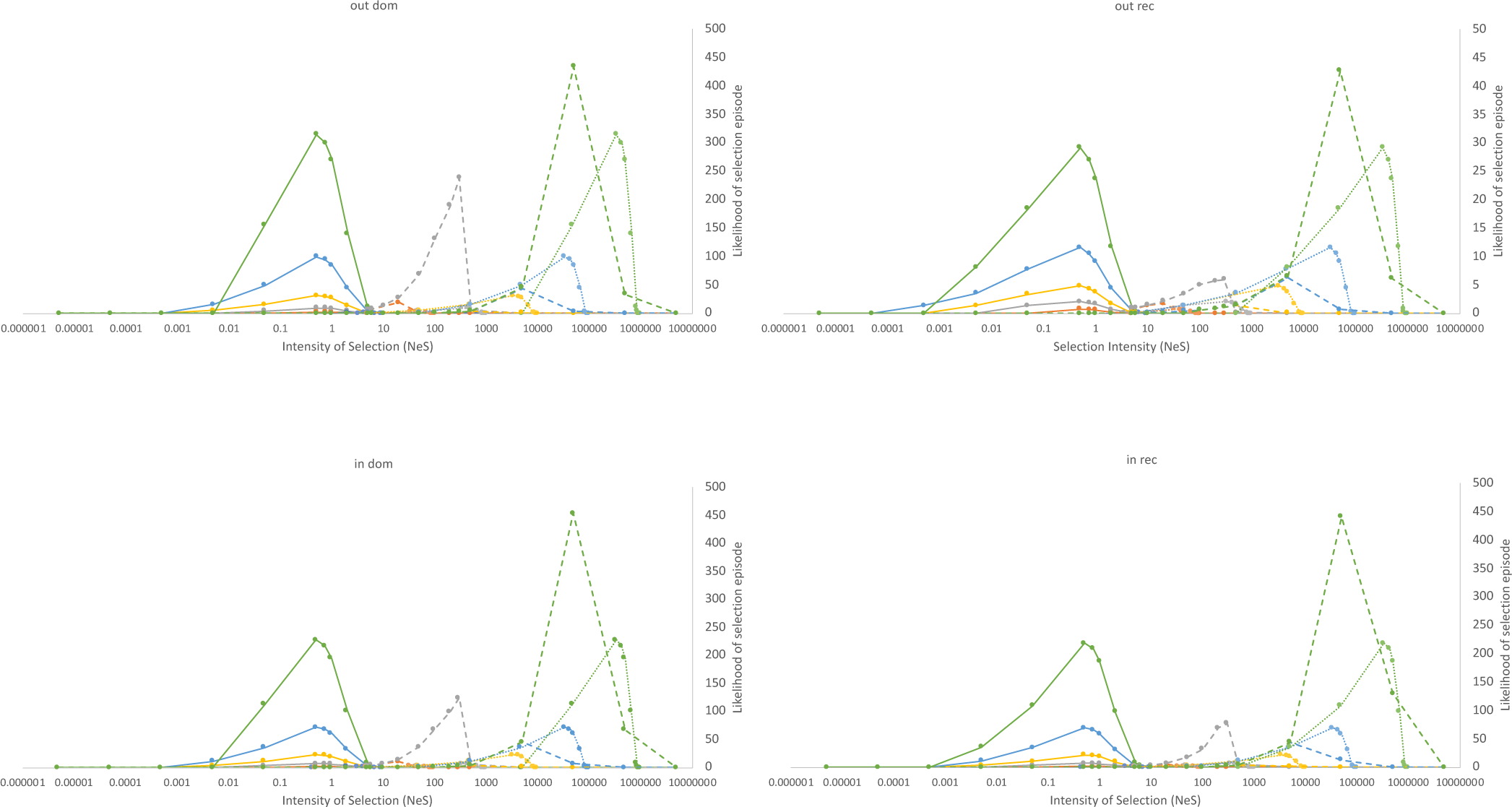
Likelihood profiles of competitive and environmental selection intensity. Likelihoods were calculated as p(threshold)*p(magnitude)*2Nµ*p(survival): the product of probabilities of a p_w_/p_m_ ratio for an associated selection intensity under an exponential distribution model of parameter λ=1, equation (11), the neutral probability of mutant frequencies becoming sufficiently high such that δp>1/2N, calculated as in Fig. S6, the probability of population survival for a given magnitude of s as calculated in (Allaby et al 2015), and the rate of mutation generation 2Nµ. Mutation rate is notionally included here as µ=0.5. Likelihoods were calculated for different population sizes as follows: blue (10), orange (100), grey (1000), yellow (10,000), light blue (100,000) and green (1000,000). Solid lines indicate competitive selection at initial intensities, dotted lines represent competitive selection at final intensities (p = 0.99), dashed lines represent environmental selection. For N >= 10,000, values for environmental selection are scaled for graphing visualization. Values are calculated under outbreeding (2% selfing) dominant (out dom), inbreeding (98% selfing) dominant (in dom), outbreeding recessive (out rec) and inbreeding recessive (in rec) conditions.

### Evidence of competitive selection in grain size evolution during domestication

We used the framework described here to investigate the patterns of selection intensity over time observed in size traits of grains in plant domestication. Using the archaeometric measurements of length, breadth and thickness of grains from nine plant species over time (Fuller *et al*. 2014), we calculated selection coefficients using haldanes based on shifting means and standard deviations over time for a given trait heritability (Kinnison and Hendry 2001), green diamonds in Fig. 2. Previously, we have found the Haldane metric approach has yielded estimates which closely match selection coefficients calculated from a Hardy-Weinberg model for shattering traits (Allaby *et al*. 2017), suggesting it is a robust approach for calculating selection strength from this data type. We examined how well the competitive selection framework described here and a scenario of shifting optimums under the standard model of stabilizing selection describes the selection coefficients over time associated with the shifts in grain size associated with stabilizing selection. Using a sliding window approach of archaeometric selection coefficients over time best fit theoretical tracks of selection coefficients were generated for each model, Fig S7. All crops show a series of episodes in which either the stabilising selection model or the competitive selection model fit best. Since the expectations of the two models are fundamentally different in that in a shifting optimum scenario is associated with diminishing selection coefficients as the new trait optimum is approached and in competitive selection coefficients increase over time, it is unsurprising that the two models are largely mutually exclusive in their fits over time. However, the fits are suggestive of a tendency for periods of competitive selection rising out of stabilizing selection episodes.

**Figure 2.**
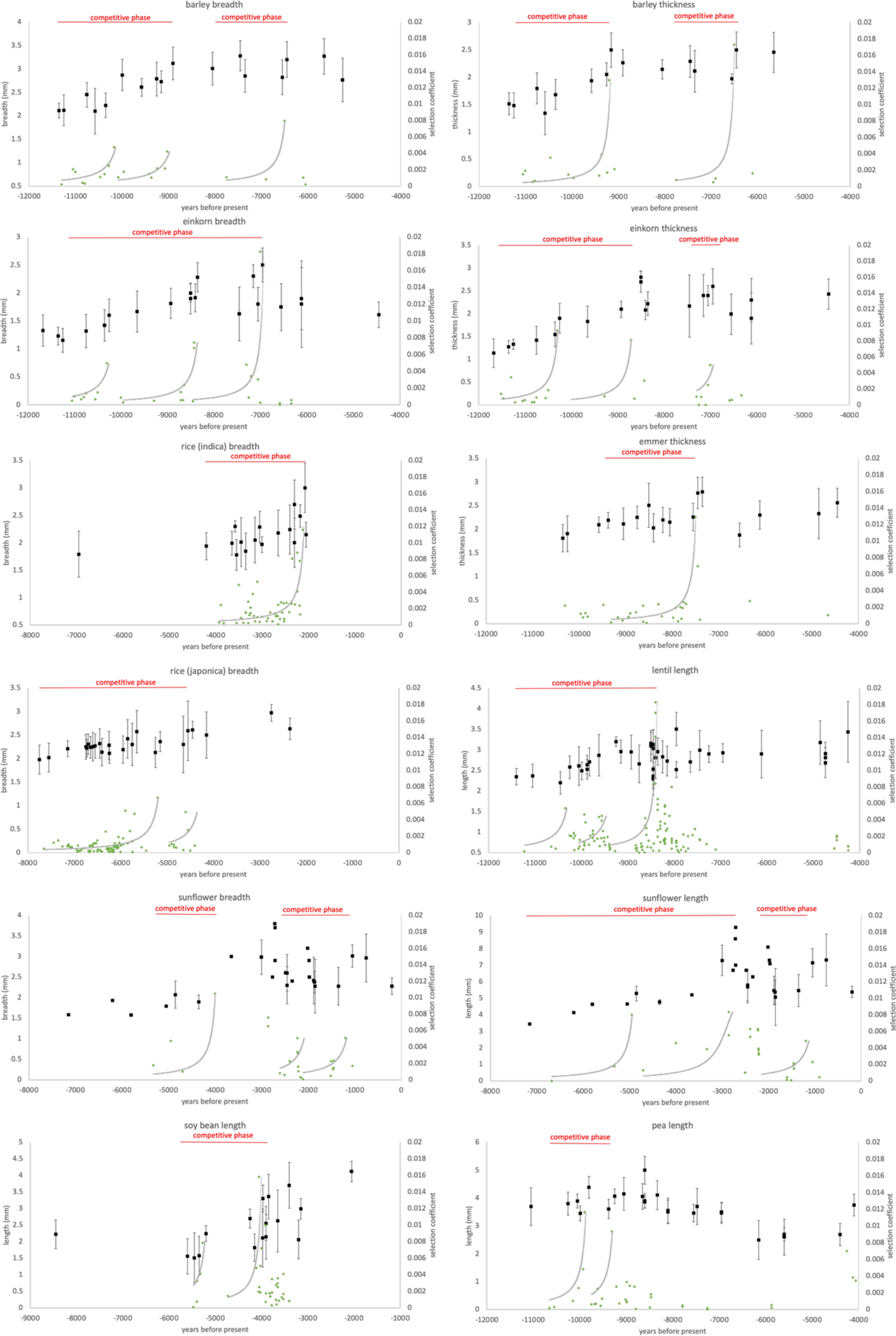
Competitive selection tracks fitted to archaeobotanical grain metric data. Grain metrics shown in black with standard deviations (axis left), data from Fuller et al 2014. Selection coefficients (green diamond, axis right) were determined by first calculating haldanes using the method of Kinnison and Hendry 2001 and converting to s following s = haldanes/H^2^ where H is the trait heritability, assumed here to be approximately 0.9 for size traits (Fuller et al 2014). Selection coefficients were binned out from high points to low points going back in time. Theoretical profiles of selection coefficients (grey lines, axis right) were determined beginning at a strength equal to s/4N of the highest selection coefficient, see methods. Profiles were fitted to archaeometric data by least squares regression of the y coordinates, r values were determined by product moment correlation coefficient pmcc on y theoretical and observed coordinates.

The time periods which fit better with a competitive selection model are characterized by trends of increasing selection coefficients. We examined 28 instances of upticks in empirical selection intensity that correlated with competitive selection episodes in more detail, improving the fit by increasing the window size and exploring a range of population sizes. Over 50% of these fits had *r* values > 0.9 and over 85% with r values > 0.75 (grey lines in Fig. 2 and Table S1). Over half the correlations showed a significantly better fit than could be achieved from fits to random upward trends (*p_null_*, Table S1), and 80% of trends that showed a strong correlation with competitive selection (*r* > 0.65) had a probability of 0.1 or less of achieving as good a fit with random trends. We therefore conclude that the majority of curves identified as a good fit to competitive selection are not over fitted, while in a few cases such as sunflower length the fit could be spurious. The latter may be resolved in the future with more archaeological data points. All instances were associated with a negative correlation to the standard model of stabilizing selection shifting optimums.

**Table 1.** Best fit parameters of theoretical competitive selection to archaeometric grain data

|  | *r* | *s* | *Nes* | *pm/pw* | Bin Dates (BP) |  | *N* | timeframe | *p* height | *p* null | standard |
| --- | --- | --- | --- | --- | --- | --- | --- | --- | --- | --- | --- |
|  |  |  |  |  | first | last |  |  |  |  | model *r* |
| barley breadth 1 | 0.83274258 | 8.04733E-06 | 0.8047328 | 1.80473928 | 11300 | 10170 | 100000 | 1130 | 0.006 | 0.0392 | -0.3750 |
| barley breadth 2 | 0.82593431 | 9.04202E-06 | 0.90420174 | 1.90420991 | 10075 | 9025 | 100000 | 1050 | 0.005 | 0.0704 | -0.3690 |
| barley breadth 3 | 0.98325575 | 8.15863E-06 | 0.40793151 | 1.40793484 | 7750 | 6500 | 50000 | 1250 | 0.019 | 0.0589 | -0.4751 |
| barley thickness 1 | 0.76449516 | 6.12979E-06 | 0.61297929 | 1.61298304 | 11050 | 9200 | 100000 | 1850 | 0.021 | 0.0361 | -0.1229 |
| barley thickness 2 | 0.99658765 | 7.34625E-06 | 0.36731245 | 1.36731514 | 7750 | 6500 | 50000 | 1250 | 0.047 | <0.001 | -0.3305 |
| einkorn breadth 1 | 0.92669622 | 1.00989E-05 | 0.50494372 | 1.50494881 | 11050 | 10300 | 50000 | 750 | 0.009 | 0.0101 | -0.2663 |
| einkorn breadth 2 | 0.75952857 | 6.07423E-06 | 0.60742346 | 1.60742715 | 9950 | 8425 | 100000 | 1525 | 0.012 | 0.0762 | -0.2656 |
| einkorn breadth 3 | 0.29709117 | 5.18241E-06 | 0.51824057 | 1.51824325 | 8450 | 7000 | 100000 | 1450 | 0.035 | 0.7295 | -0.3055 |
| einkorn thickness 1 | 0.88384627 | 9.09309E-06 | 0.90930918 | 1.90931744 | 11512 | 10300 | 100000 | 1212 | 0.01 | 0.0060 | -0.0468 |
| einkorn thickness 2 | 0.99631575 | 7.1627E-06 | 0.35813512 | 1.35813768 | 10000 | 8712 | 50000 | 1288 | 0.022 | <0.001 | -0.5221 |
| einkorn thickness 3 | 0.94067196 | 6.05017E-06 | 0.60501711 | 1.60502077 | 7300 | 7000 | 100000 | 300 | 0.008 | 0.0251 | -0.5387 |
| emmer thickness 1 | 0.94465145 | 8.12901E-06 | 0.81290079 | 1.8129074 | 9312 | 7500 | 100000 | 1812 | 0.015 | <0.001 | -0.1439 |
| rice (indica) breadth 1 | 0.80466945 | 5.11564E-06 | 0.51156442 | 1.51156704 | 3925 | 2125 | 100000 | 1800 | 0.023 | <0.001 | -0.1141 |
| rice (japonica) breadth 1 | 0.67239719 | 2.16677E-05 | 0.21667677 | 1.21668147 | 7650 | 5200 | 10000 | 2450 | 0.029 | <0.001 | -0.0422 |
| rice (japonica) breadth 2 | 0.65032522 | 1.84918E-05 | 0.18491785 | 1.18492127 | 4950 | 4600 | 10000 | 350 | 0.026 | 0.0941 | -0.1409 |
| lentil length 1 | 0.93984652 | 6.05421E-06 | 0.60542145 | 1.60542512 | 11225 | 10350 | 100000 | 875 | 0.009 | 0.1093 | -0.4127 |
| lentil length 2 | 0.45014131 | 9.04643E-06 | 0.90464278 | 1.90465096 | 10062 | 9562 | 100000 | 500 | 0.005 | 0.2052 | -0.2747 |
| lentil length 3 | 0.84050712 | 6.18286E-06 | 0.61828609 | 1.61828991 | 9385 | 8400 | 100000 | 985 | 0.029 | 0.0010 | -0.1463 |
| pea length 1 | 0.94233761 | 5.11646E-06 | 0.51164627 | 1.51164889 | 10650 | 9895 | 100000 | 755 | 0.022 | 0.0262 | -0.5338 |
| pea length 2 | 0.96610421 | 6.09332E-06 | 0.6093318 | 1.60933551 | 9750 | 9312 | 100000 | 438 | 0.015 | 0.0020 | -0.2292 |
| sunflower length 1 | 0.98754599 | 6.07981E-06 | 0.6079807 | 1.60798439 | 6675 | 4950 | 100000 | 1725 | 0.013 | 0.1597 | -1.0000 |
| sunflower length 2 | 0.82637007 | 9.33817E-06 | 0.00933817 | 1.00933825 | 4700 | 2855 | 1000 | 1845 | 0.896 | 0.2298 | -0.7477 |
| sunflower length 3 | 0.92894554 | 1.7484E-05 | 0.17484025 | 1.17484331 | 2155 | 1200 | 10000 | 955 | 0.027 | 0.1081 | -0.5499 |
| sunflower breadth 1 | 0.92332224 | 6.10484E-06 | 0.61048379 | 1.61048752 | 5325 | 4000 | 100000 | 1325 | 0.017 | 0.2515 | -0.4237 |
| sunflower breadth 2 | 0.81764058 | 1.50505E-05 | 0.01505051 | 1.01505073 | 2607 | 2215 | 1000 | 392 | 0.328 | 0.0270 | -0.4531 |
| sunflower breadth 3 | 0.93326734 | 9.05106E-06 | 0.90510587 | 1.90511406 | 2110 | 1200 | 100000 | 910 | 0.005 | 0.0461 | -0.4763 |
| soybean length 1 | 0.96449128 | 7.07858E-06 | 0.70785833 | 1.70786334 | 5475 | 5275 | 100000 | 200 | 0.011 | 0.0531 | -0.5926 |
| soybean length 2 | 0.54628783 | 1.11584E-05 | 1.11583602 | 2.11584847 | 4725 | 4060 | 100000 | 665 | 0.014 | 0.7487 | -0.4561 |

The close correlation between the selection dynamics of complex size traits and the competitive model which considers the fate of single adaptive alleles, suggests that in most cases during the intensification of selection, single alleles of large effect are predominately responsible for shifts in size traits during these episodes. In its current form the competitive selection framework does not consider multiple alleles contributing to a trait, so we cannot entirely exclude this as a possibility, although a multigenic framework would be highly complex with dynamic values of *p_m_* which would not be likely to follow the framework outlined here. In some cases concurrent competitive episodes were identified for different traits in the same species, such as in the case of barley breadth and thickness. It is possible that multiple alleles hitchhiked together each affecting one trait, or that a single allele of large effect was pleiotropic in its trait effects. In other cases, such as the first two episodes in both sunflower breadth and length, they appear more staggered, suggesting a complex sequence of episodes that we can take to suggest different alleles being selected sequentially. The frequency of *p* at the height of the competitive episode trend (*p* height, Table S1) was generally found to be quite low, usually between 1-5%, Table S1. It is unlikely this represents the final frequency that a putative adaptive allele reached. Since the log phase of frequency increase is short under this model at these intensities, typically less than 100 generations between 0.02 and 0.95, the density of archaeological sampling is unlikely to recover the latter stages of a fixation episode. We cannot therefore conclude from these data whether these signals are associated with allele fixations under selection.

In most cases a model is supported of multiple competitive episodes for each crop. The fitted selection intensities on archaeometric data match remarkably closely to the theoretical expected intensities described in the model, Fig. 3, indicating adaptive advantages that range from a 0.9-116% improvement in resource acquisition with a mode of 65%. Given the close correlations of the pattern of selection intensities to the empirical archaeometric data, and the close match to the expected theoretical intensities, we conclude that these nine crop species most likely underwent competitive selection episodes which often appear to have emerged during periods of shifting optimums in stabilizing selection.

**Figure 3.**
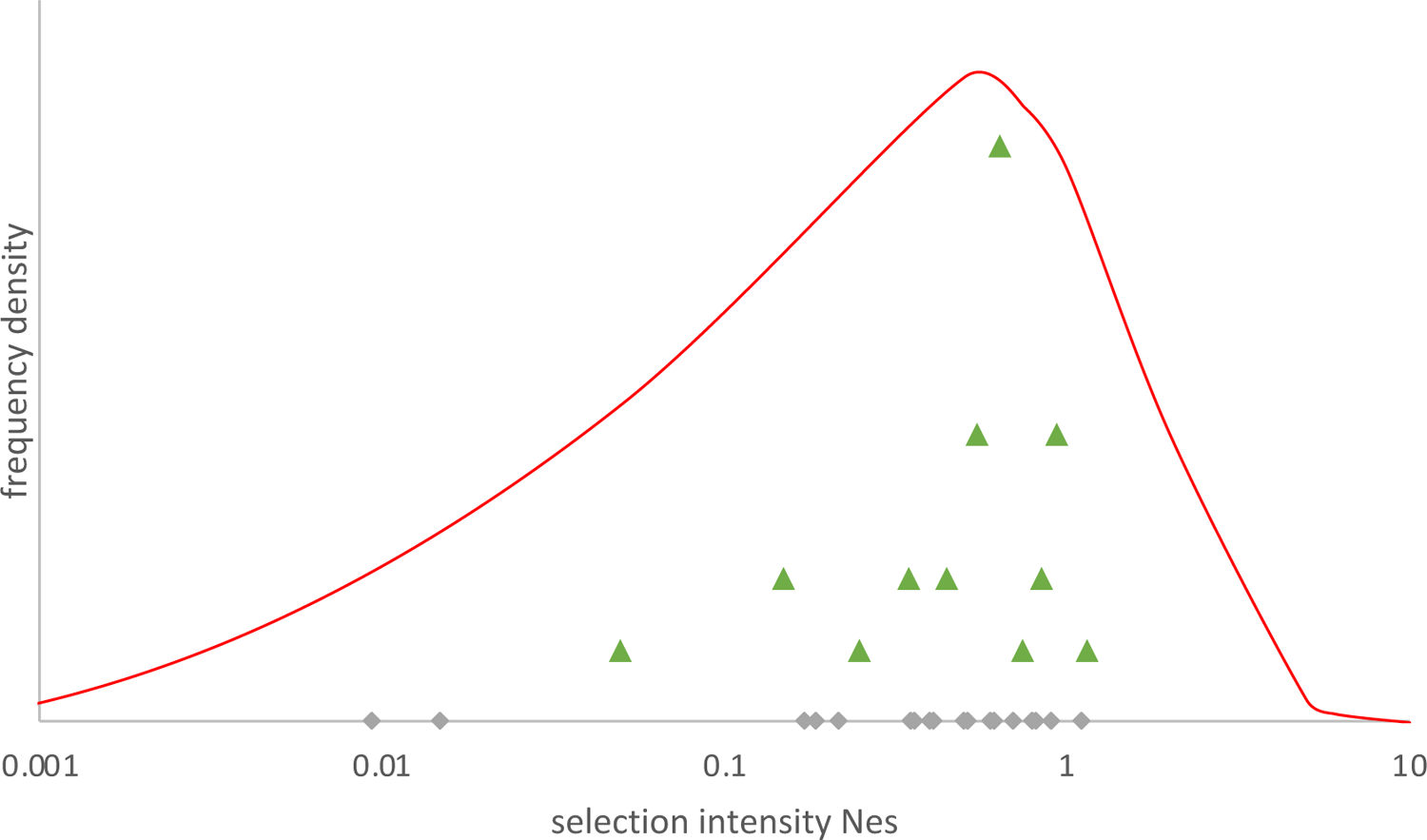
Best fit competitive selection intensities on archaeometric grain data and theoretical intensity likelihood profile for inbreeding dominant crops. Red line depicts the frequency density based on N=100,000 likelihood profile for inbreeding dominant plants as calculated in Fig. 2 (note that inbreeding recessive, out breeding dominant and recessive do not differ greatly when scaled). Grey diamonds indicate selection intensity values determined in Table S1, green triangles indicate interval frequencies of determined selection intensities from Table S1.

## DISCUSSION

Complex traits such as size are well known to be under stabilizing selection, which has a well established model framework (Barton 1986). A prediction of this model is that as traits approach their optimal value the selection pressure becomes less, as selection strength is a function of the difference between the trait value and the trait optimum. In part, this system is derived from the observation that under selective breeding the rate of crop trait change diminishes over time as improvements become constrained by the pleiotropic effects of other traits (Dudley and Lambert 2004, Johnson and Barton 2005). Consequently, the observation of increasing selection intensity over time of complex size traits from archaeometric data appears anomalous to stabilizing selection as it is currently understood.

Cultivation led in almost all cases of crops to an increase in grain size dimensions over several millennia (Fuller *et al*. 2014). The new environment of cultivation may have heralded a new set of optimums for trait dimensions which encouraged a shift in size. In particular it is argued that by provision of stable resources through cultivation the optimal strategy for plants shifted from nutrient conservation to nutrient acquisition (Fuller and Stevens 2017, Miller *et al*. 2015). Models of stabilizing selection suggest that it is alleles of small effect which are most likely to respond first to shifts in optima, partly because they are expected to occur at intermediate frequencies rather than either rare or fixed in the case of large effect alleles (De Vladar and Barton 2014). Furthermore, selective sweeps are expected to be rare (Thornton 2019), and where alleles of large effect are involved, small effect alleles are likely to replace them in the course of adaptation (Chevin and Hospital 2008). Under these circumstances, we would expect trait adjustment to follow a pattern of decelerating selection pressure. The pattern we observe in the archaeological record of increasing intensity repeatedly in many crops is suggestive that an alternative process to the standard model of stabilizing selection is involved.

If a new trait optimum is greatly removed from the current trait mean, outside the range of trait effects in the current genetic variation, it may be that large jumps may not be within evolutionary reach (Jain and Stephan 2017). Escape from such local optima traps can be achieved through new mutations of large effect which can launch the phenotype to a new part of the adaptive landscape (Jain and Stephan 2017). The patterns here suggest that such leaps may have occurred during shifting optimum episodes initiated by the move to a cultivated environment, reflecting a switch in environmentally driven selection to competitive driven selection. This fits the notion of alleles of large effect enabling crops to capitalize on resource availability through larger grain size at the expense of neighbour resource availability during shifting optimum episodes, which in turn could have led to robust growth and seedling competition, for instance shading wild type plants. The fit with the competitive model presented here suggests simple single gene driven changes, and although we cannot rule out multigenic systems on the evidence here we do not expect they would follow the same trajectories because values of *p_m_* would necessarily be dynamic with time as various combinations of contributing alleles were assembled in individuals over time. It may be the case that such alleles became fixed, or, numerous other loci of lesser effect could have come into play in later stages (Chevin and Hospital 2008). Archaeometric evidence is unlikely to capture the later stages of competitive episodes, which are necessarily brief, but it is an expectation of the model that such alleles of large effect would have been fixed. It is likely that on fixation of alleles of large effect there would still have been a range alleles of small effect in the population, and so a return to an equilibrium of stabilizing selection. After competitive episodes, trait values appear to slow down in their rate of increase, and in some instances we see reducing trends in selection intensities, which appears to be reflective of a standard stabilizing selection model as traits appear to reach pleiotropically constrained new optima.

## CONCLUSION

The cost consequences of adaptive mutations in the absence of environmental deterioration have largely been overlooked since the conception of the substitution load. As was initially suggested (Brues 1964), we find that the initial impact on the population is very low, but the cost consequence for the wild type becomes significant with time and its properties as described here are detectable in time series data from which selection coefficients can be derived. Consequently, the approaches outlined here are applicable to archaeogenomic studies of ancient DNA over time. Our study suggests that in crop domestication competitive selection was a widespread occurrence and provides a new complexity to modelling the evolution of complex traits. It remains to be seen how widespread such selection might have been in wild populations.

## Author contributions

RGA designed and performed the analysis, CJS and DQF provided archaeometric data, RGA wrote the manuscript with input from CJS and DQF.

## Acknowledgements

The authors would like to thank Logan Kistler, Becky Cribdon and Hsiao-Lei Liu for useful discussions in developing this manuscript.

## Data availability

All archaeometric data are published and custom scripts used in analyses are available from https://warwick.ac.uk/fac/sci/lifesci/research/archaeobotany/downloads/Competitive_selection_scripts

## Supplementary Information

**Figure S1.**
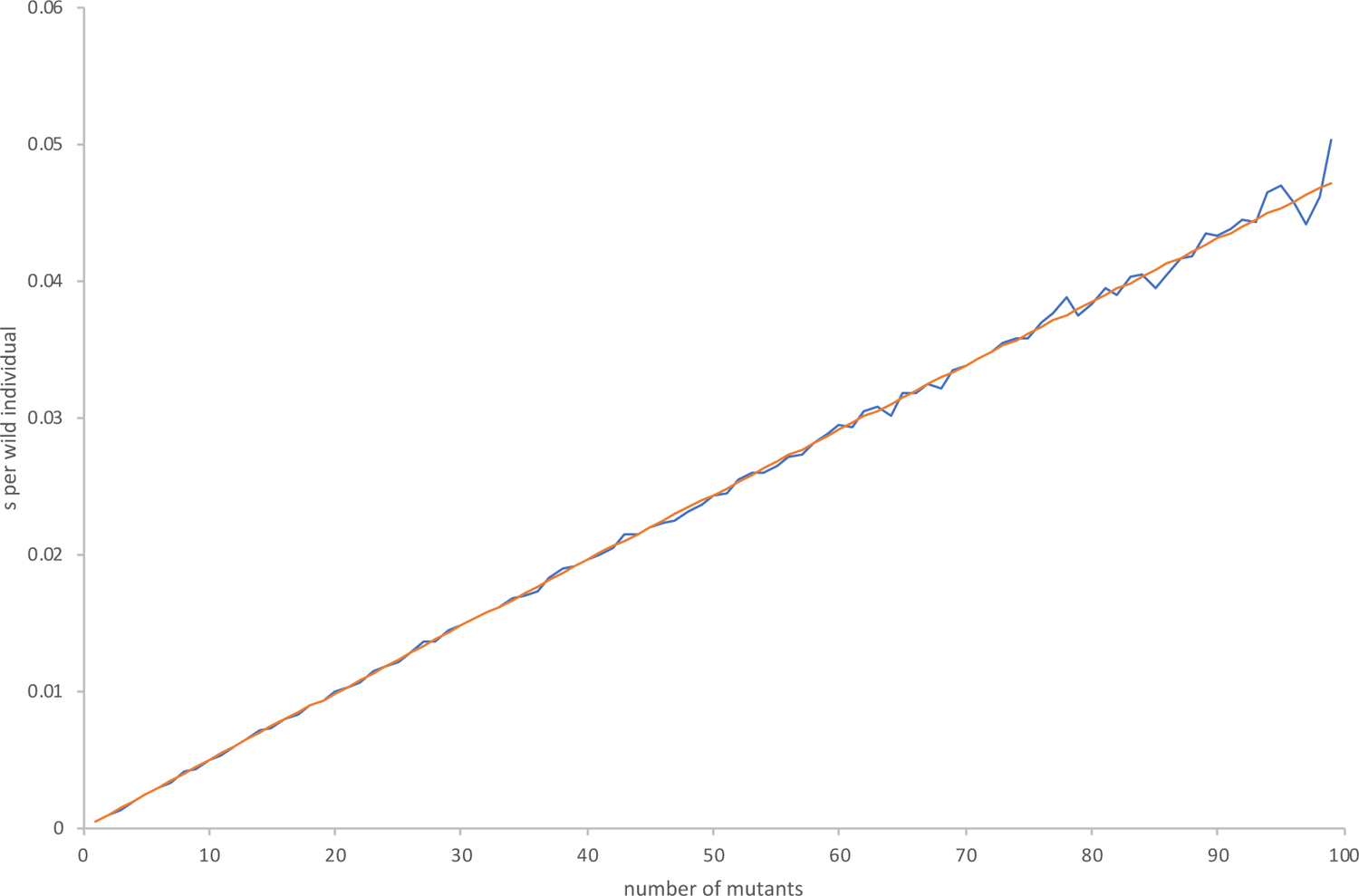
Observed and theoretical proportional loss (s) of resource in wild type under conditions of benign mutant selection. Observed values (blue) generated assigning values of 0.21 and 0.2 to p_m_ and p_w_ respectively in a population size of 100 individuals with resource size 1000 and 10,000 replications. Individuals were randomly selected and then deemed successful in resource acquisition with a probability of p_m_ or p_w_ depending on whether they were wild type or mutant individuals. Sampling continued until all resource had been allocated. Model available as script benign_profile_of_s.pl from (website). Theoretical values (orange) calculated from equation (6) using the same values of p_m_ and p_w_.

**Figure S2.**
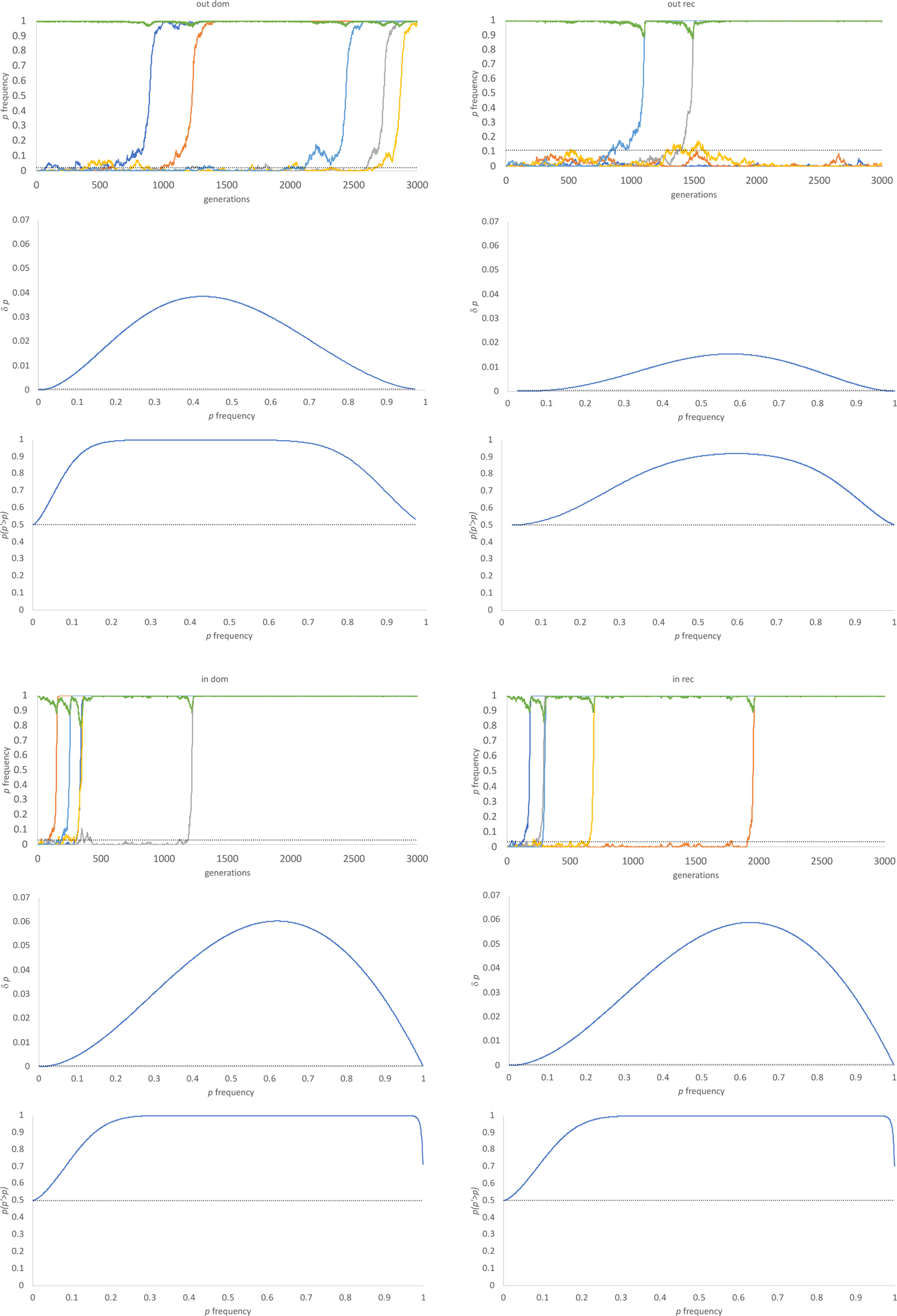
Competitive selection frequency dependent thresholds. Top panels: competitive selection simulations at 5 loci, for s= 0.0005 and N = 1000. Frequency of selected mutant (p) denoted by colours (dark blue, orange, grey, yellow and light blue respectively), population size N shown in green. Threshold frequencies of p at which δp>1/2N shown by dotted line. Middle panels: Values of δp at given frequencies of p calculated using equation (8) using p dependent values of s calculated in equation (6), for which values of N_m_ and N_w_ were calculated in equations (12) and (13). Threshold at which δp = 1/2N shown by dotted line. Bottom panels: Probabilities of p’>p. Values of p’ = p + δp. Normal approximation to the binomial distribution used in which µ = 2Np’ and σ = (2Np’(1-p’)^0.5^ and cumulative probabilities calculated for x>p. Pure genetic drift levels are shown by dotted line where µ = p. Simulations carried out under outbreeding (2% selfing) dominant (out dom), inbreeding (98% selfing) dominant (in dom), outbreeding recessive (out rec) and inbreeding recessive (in rec) conditions.

**Figure S3.**
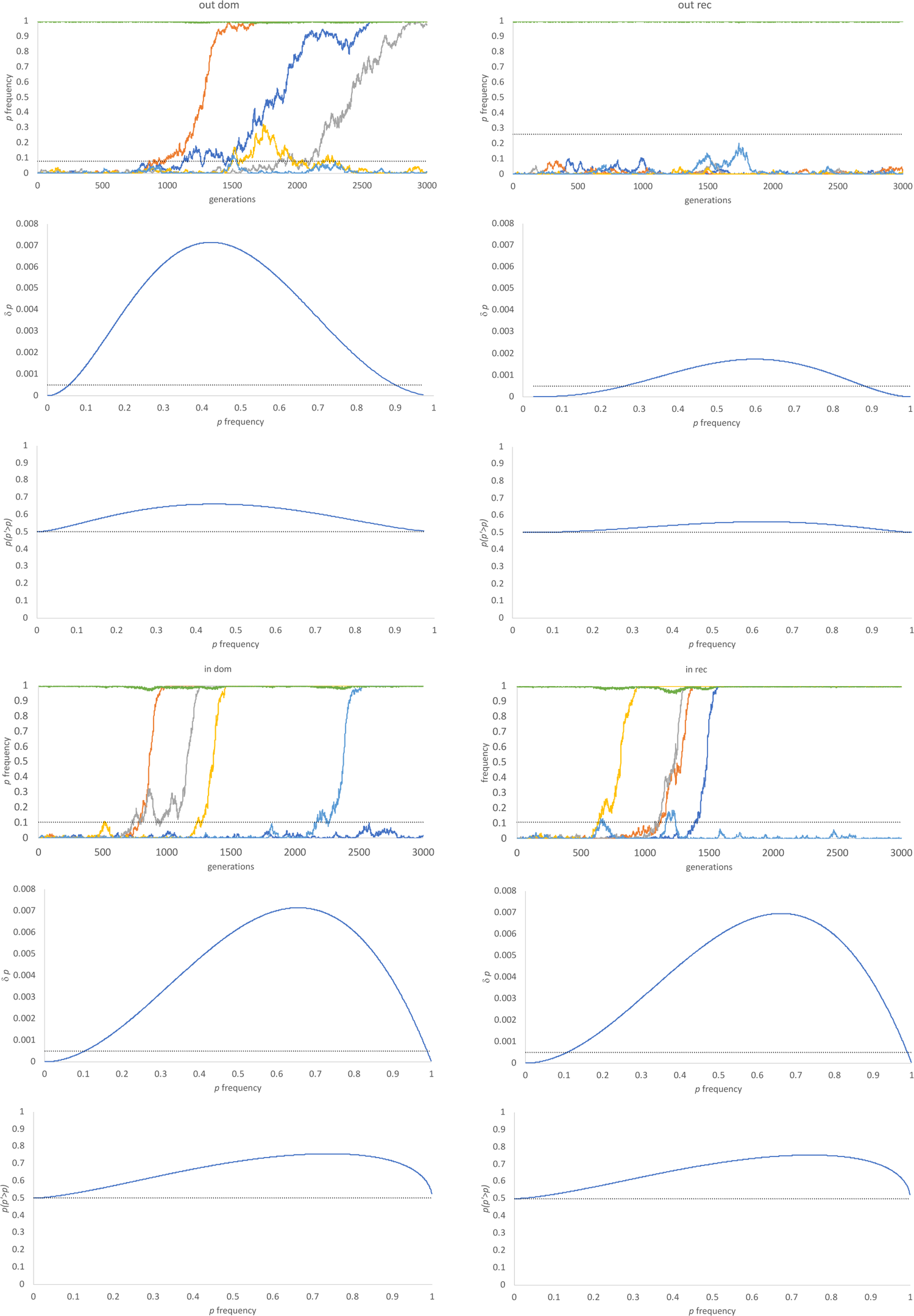
Competitive selection frequency dependent thresholds. Top panels: competitive selection simulations at 5 loci, for s= 0.00005 and N = 1000. Frequency of selected mutant (p) denoted by colours (dark blue, orange, grey, yellow and light blue respectively), population size N shown in green. Threshold frequencies of p at which δp>1/2N shown by dotted line. Middle panels: Values of δp at given frequencies of p calculated using equation (8) using p dependent values of s calculated in equation (6), for which values of N_m_ and N_w_ were calculated in equations (12) and (13). Threshold at which δp = 1/2N shown by dotted line. Bottom panels: Probabilities of p’>p. Values of p’ = p + δp. Normal approximation to the binomial distribution used in which µ = 2Np’ and σ = (2Np’(1-p’)^0.5^ and cumulative probabilities calculated for x>p. Pure genetic drift levels are shown by dotted line where µ = p. Simulations carried out under outbreeding (2% selfing) dominant (out dom), inbreeding (98% selfing) dominant (in dom), outbreeding recessive (out rec) and inbreeding recessive (in rec) conditions.

**Figure 4.**
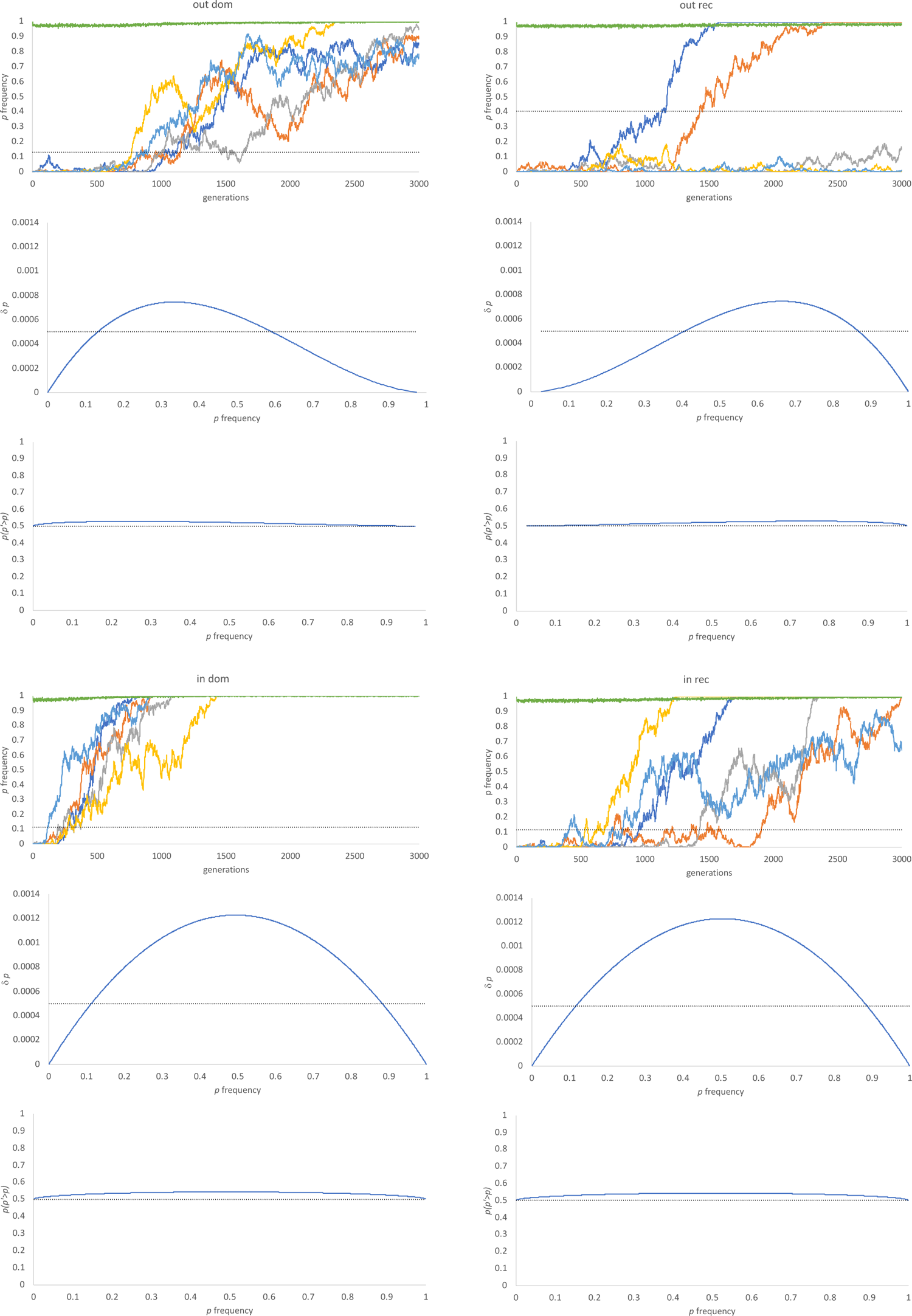
Environmental selection frequency dependent thresholds. Top panels: environmental selection simulations at 5 loci, for s= 0.005 and N = 1000. Frequency of selected mutant (p) denoted by colours (dark blue, orange, grey, yellow and light blue respectively), population size N shown in green. Threshold frequencies of p at which δp>1/2N shown by dotted line. Middle panels: Values of δp at given frequencies of p calculated using equation (8) using p dependent values of s calculated in equation (6). Threshold at which δp = 1/2N shown by dotted line. Bottom panels: Probabilities of p’>p. Values of p’ = p + δp. Normal approximation to the binomial distribution used in which µ = 2Np’ and σ = (2Np’(1-p’)^0.5^ and cumulative probabilities calculated for x>p. Pure genetic drift levels are shown by dotted line where µ = p. Simulations carried out under outbreeding (2% selfing) dominant (out dom), inbreeding (98% selfing) dominant (in dom), outbreeding recessive (out rec) and inbreeding recessive (in rec) conditions.

**Figure S5.**
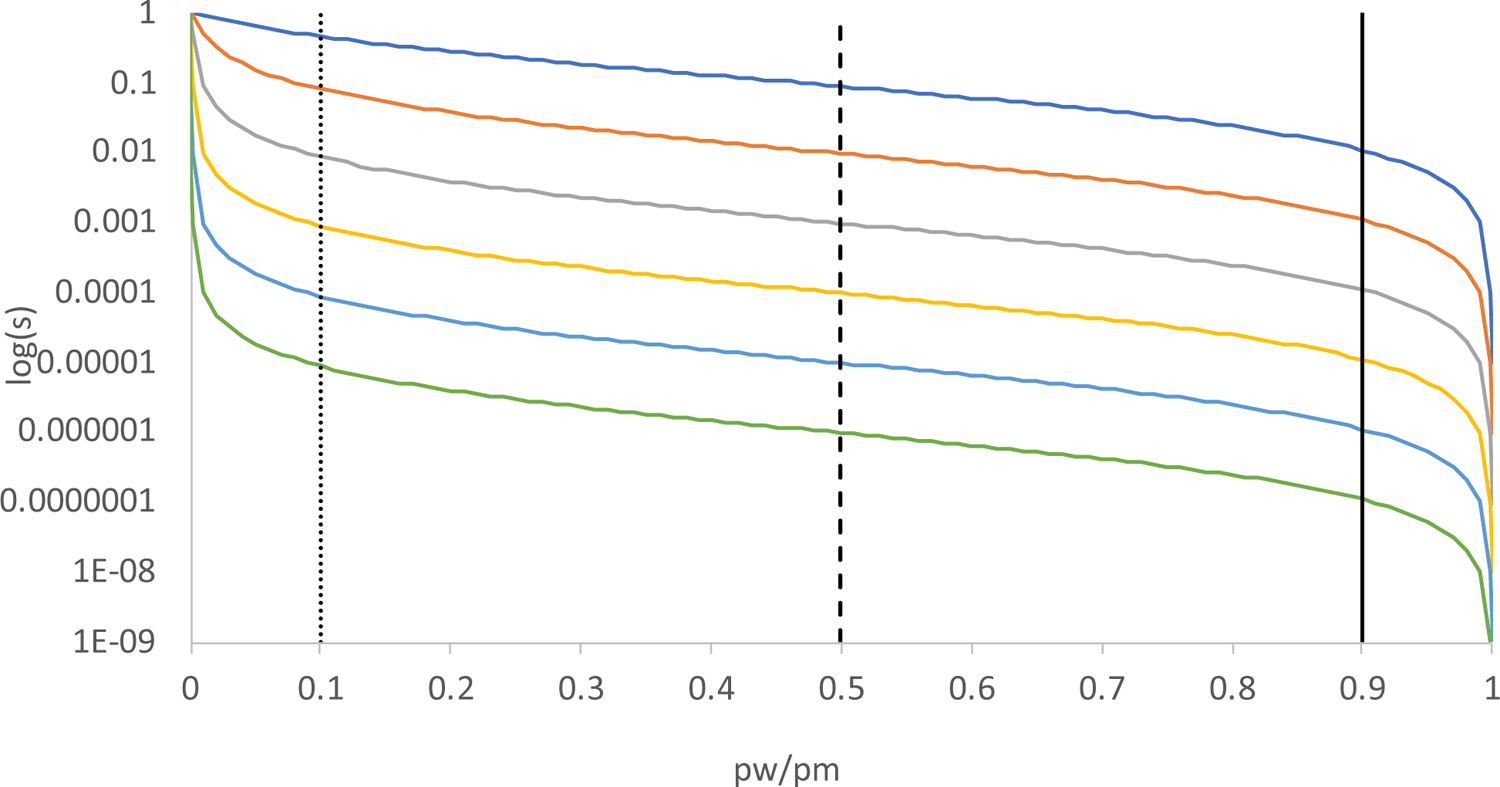
The relationship between pw/pm and s for given population sizes. Values of s were derived from equation (6) for given p_w_/p_m_ values for the following population sizes: blue (10), orange (100), grey (1000), yellow (10,000), light blue (100,000) and green (1000,000). Vertical bars denote selection intensities of 0.1 (solid), 1 (dashed) and 10 (dotted).

**Figure S6.**
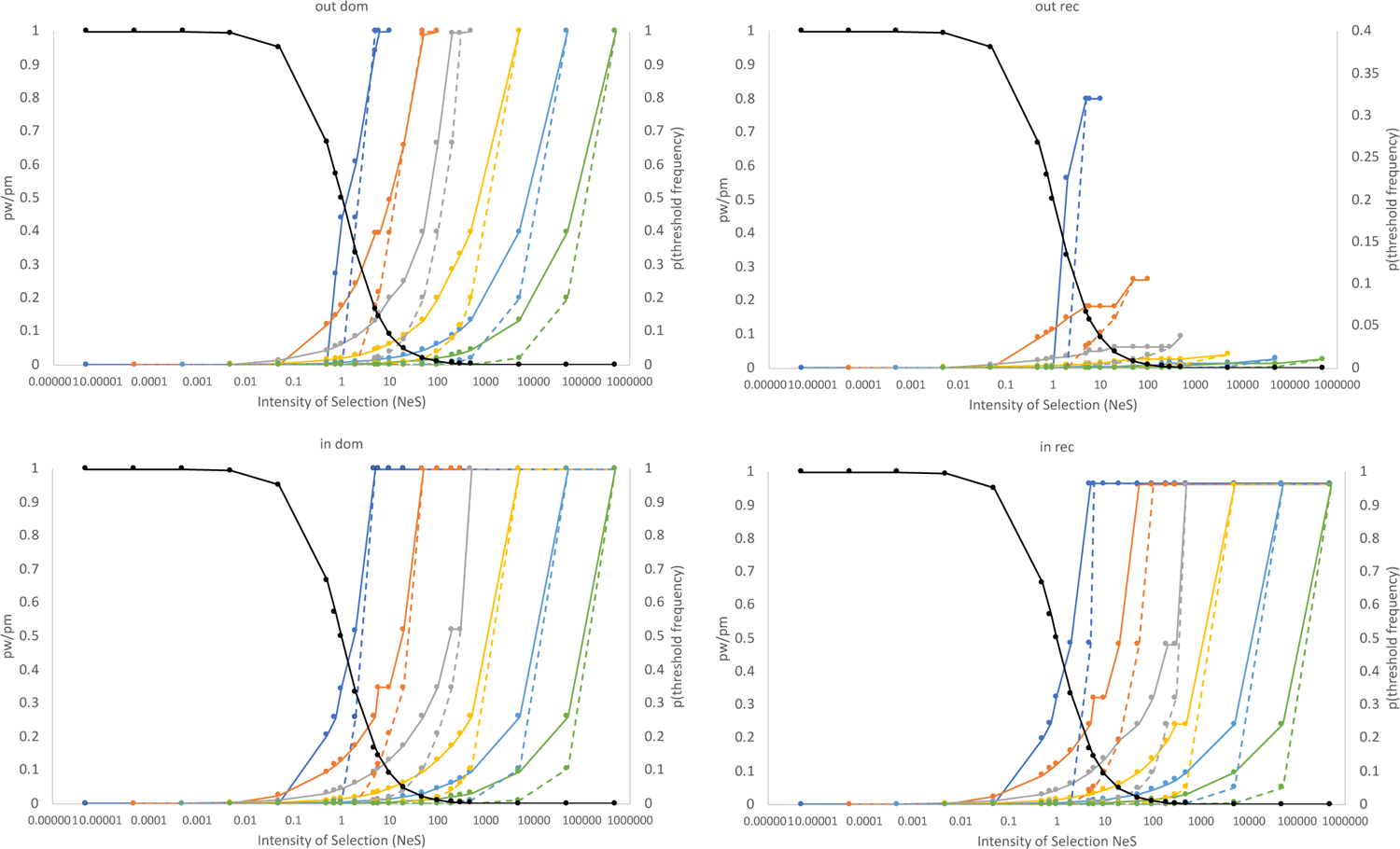
Probability of reaching drift/selection threshold frequencies for given selection intensities and their associated p_w_/p_m_ ratios. The p_w_/p_m_ ratios shown in dark black. Probabilities of reaching threshold frequencies through drift were calculated by first determining the number of individuals at which δp > 1/2N using equations and (10) to determine p values for given N_m_ values, equation (6) to determine the frequency dependent values of s and equation (8) to determine δp. The neutral probability of an allele reaching that frequency is given as 1/f_(threshold)_. Probabilities were calculated for population sizes of blue (10), orange (100), grey (1000), yellow (10,000), light blue (100,000) and green (1000,000), solid lines represent competitive selection probabilities and dashed lines represent the corresponding environmental selection probabilities. Values are calculated under outbreeding (2% selfing) dominant (out dom), inbreeding (98% selfing) dominant (in dom), outbreeding recessive (out rec) and inbreeding recessive (in rec) conditions.

**Figure S7.**
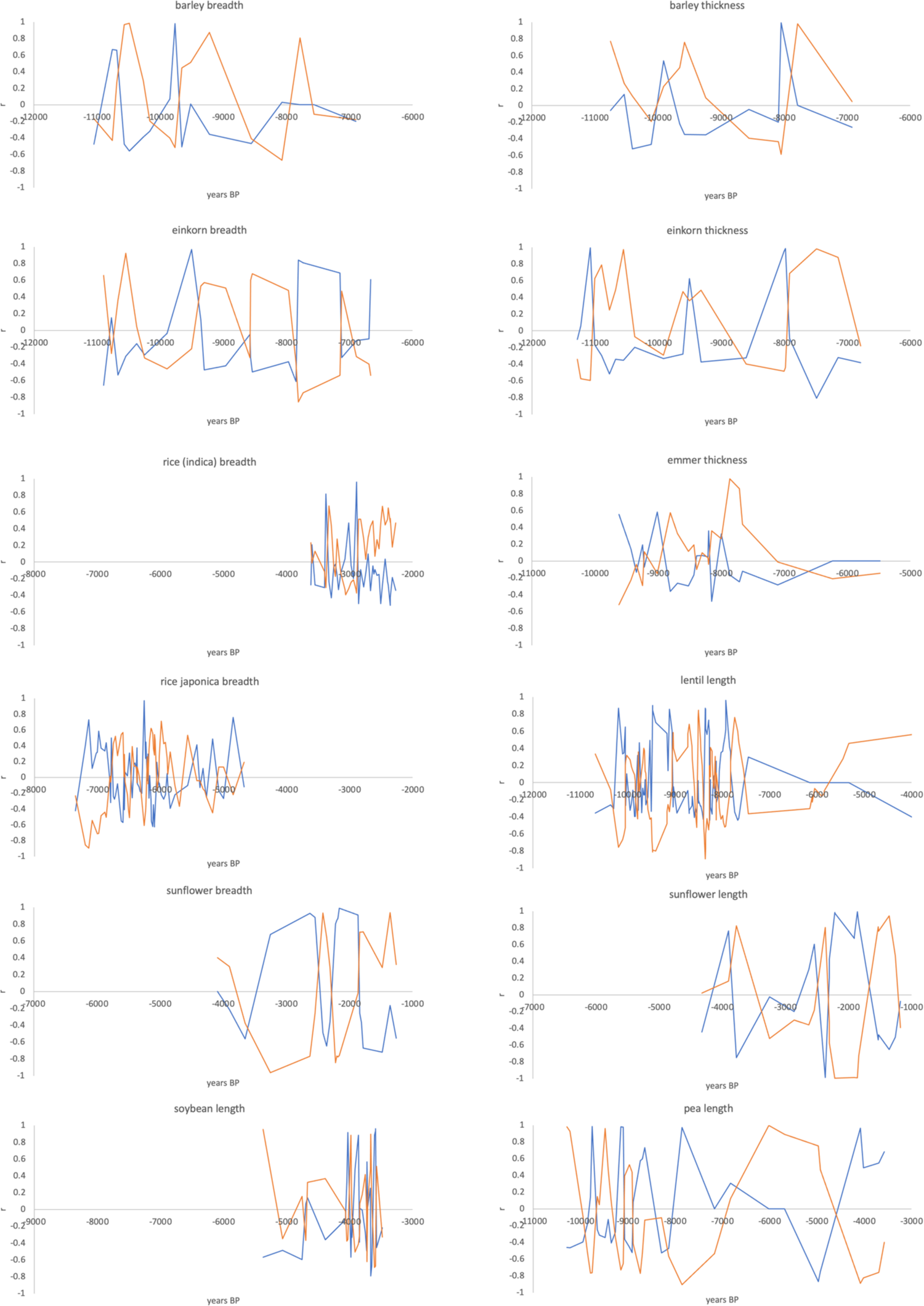
Competitive and shifting optimum stabilizing selection model fit to archaeometric based selection coefficients. Graphs show correlation coefficient values for best fit selection tracks to real archaeometric data for competitive (orange) and shifting optimum stabilizing selection (blue). See methods for details of fitting and track generation.

## Literature Cited

1. Allaby RG, Kitchen JL, Fuller DQ (2015) Surprisingly low limits of selection in plant domestication. Evolutionary Bioinformatics 11:(S2) 41–51.

2. Allaby, R. G., Stevens, S., Lucas, L., Maeda, O., & Fuller, D. Q. (2017). Geographic mosaics and changing rates of cereal domestication. Philosophical Transactions of the Royal Society B, 372, 20160429.

3. Barton NH (1986) The maintenance of polygenic variation through a balance between mutation and stabilizing selection *Genet*. Res. Camb. 47:209–216.

4. Barton NH (1995) Linkage and the limits to natural selection. Genetics 140:821–841

5. Brues AM (1964) The cost of evolution versus the cost of not evolving. Evolution 18:379-383.

6. Brues AM (1969) Genetic load and its varieties. Science 164:1130–1136.

7. Chevin L.M., Hospital F (2008) Selective sweep at a quantitative trait locus in the presence of background genetic variation. Genetics 180:1645–1660.

8. De Vladar HP, Barton N (2014) Stability and response of polygenic traits to stabilizing selection and mutation. Genetics 197:749–767.

9. Dudley JW, Lambert RJ (2004) 100 generations of selection for oil and protein in corn. Plant Breeding Rev. 24:79–110.

10. Fuller, DQ and Stevens, CJ (2017) Open for Competition: Domesticates, Parasitic Domesticoids and the Agricultural Niche. Archaeology International 20: 110–121.

11. Haldane JBS. (1957) The cost of selection. J Genet. 55:511–24.

12. Fuller DQ, Denham T, Arroyo-Kalin M, Lucas L, Stephens C, Qin L, Allaby RG, Purugganan MD (2014) Convergent evolution and parallelism in plant domestication revealed by an expanding archaeological record. Proc. Natl. Acad. Sci. U.S.A. 111:6147–6152.

13. Fuller, D.Q. and Stevens C.J. (2019) The making of the botanical battleground: domestication and the origins of the World’s weed floras. In Far from the Hearth: Essays in Honour of Martin K. Jones (edited by E. Lightfoot, X. Liu, and D. Q. Fuller). Cambridge: McDonald Institute of Archaeology. Pp. 9–21.

14. Gillespie, JH (1983) A simple stochastic gene substitution model. Theor. Popul. Biol. 23: 202–215.

15. Gillespie, JH (1984) Molecular evolution over the mutational landscape. Evolution 38: 1116– 1129.

16. Jain K, Stephan W (2017) Rapid adaptation of a polygenic trait after a sudden environmental shift. Genetics 206:389–406.

17. Johnson T, Barton N (2005) Theoretical models of selection and mutation on quantitative traits. Phil. Trans R. Soc. B. 360:1411–1425.

18. Kinnison MT, Hendry AP (2001) The pace of modern life II: From rates of contemporary microevolution to pattern and process. Genetica 112-113(1):145–164.

19. Maynard-Smith J.(1968) “Haldane’s dilemma” and the rate of evolution. Nature 219:1114–6

20. Milla R, Osborne CP, Turcotte MM, Violle C (2015) Plant domestication seen through an ecological lens. Trends Ecol. Evol. 30(8):463-469.

21. Orr, HA (2003) The distribution of fitness effects among beneficial mutations. Genetics 163: 1519–1526.

22. Pavlidis P, Metzler D, Stephan W (2012) Selective sweeps in multilocus models of quantitative traits. Genetics 192:225–239.

23. Sved, JA (1968) Possible rates of gene substitution in evolution. Amer. Natur. 102:283–293.

24. Thornton K (2019) Polygenic adaptation to an environmental shift: temporal dynamics of variation under Gaussian stabilizing selection and additive effects on a single trait. Genetics 213:1513–1530.

25. Van Valen L (1963) Haldane’s dilemma, evolutionary rates, and heterosis. Amer. Natur. 97:185–190.

